# Antisense pairing and SNORD13 structure guide RNA cytidine acetylation

**DOI:** 10.1101/2022.05.12.491732

**Authors:** Supuni Thalalla Gamage, Marie-Line Bortolin-Cavaillé, Courtney Link, Keri Bryson, Aldema Sas-Chen, Schraga Schwartz, Jérôme Cavaillé, Jordan L. Meier

**Affiliations:** Chemical Biology Laboratory, National Cancer Institute, Frederick, Maryland 21702, United States; Molecular, Cellular and Developmental Biology unit (MCD), Centre de Biologie Integrative (CBI), University of Toulouse; UPS; CNRS; 118 route de Narbonne, 31062 Toulouse, France; The Shmunis School of Biomedicine and Cancer Research, The George S. Wise Faculty of Life Sciences, Tel Aviv University, Tel Aviv, Israel; Department of Molecular Genetics, Weizmann Institute of Science, Rehovot, Israel

## Abstract

N4-acetylcytidine (ac^4^C) is an RNA nucleobase found in all domains of life. Establishment of ac^4^C in helix 45 (h45) of human 18S ribosomal RNA (rRNA) requires the combined activity of the acetyltransferase NAT10 and the box C/D snoRNA SNORD13. However, the molecular mechanisms governing RNA-guided nucleobase acetylation in humans remain unexplored. Here we report two assays that enable the study of SNORD13-dependent RNA acetylation in human cells. First, we demonstrate that ectopic expression of SNORD13 rescues h45 in a SNORD13 knockout cell line. Next, we show mutant snoRNAs can be used in combination with nucleotide resolution ac^4^C sequencing to define structure and sequence elements critical for SNORD13 function. Finally, we develop a second method that reports on the substrate specificity of endogenous NAT10-SNORD13 via mutational analysis of an ectopically-expressed pre-rRNA substrate. By combining mutational analysis of these reconstituted systems with nucleotide resolution ac^4^C sequencing, our studies reveal plasticity in the molecular determinants underlying RNA-guided cytidine acetylation that is distinct from deposition of other well-studied rRNA modifications (e.g. pseudouridine). Overall, our studies provide a new approach to reconstitute RNA-guided cytidine acetylation in human cells as well as nucleotide resolution insights into the mechanisms governing this process.

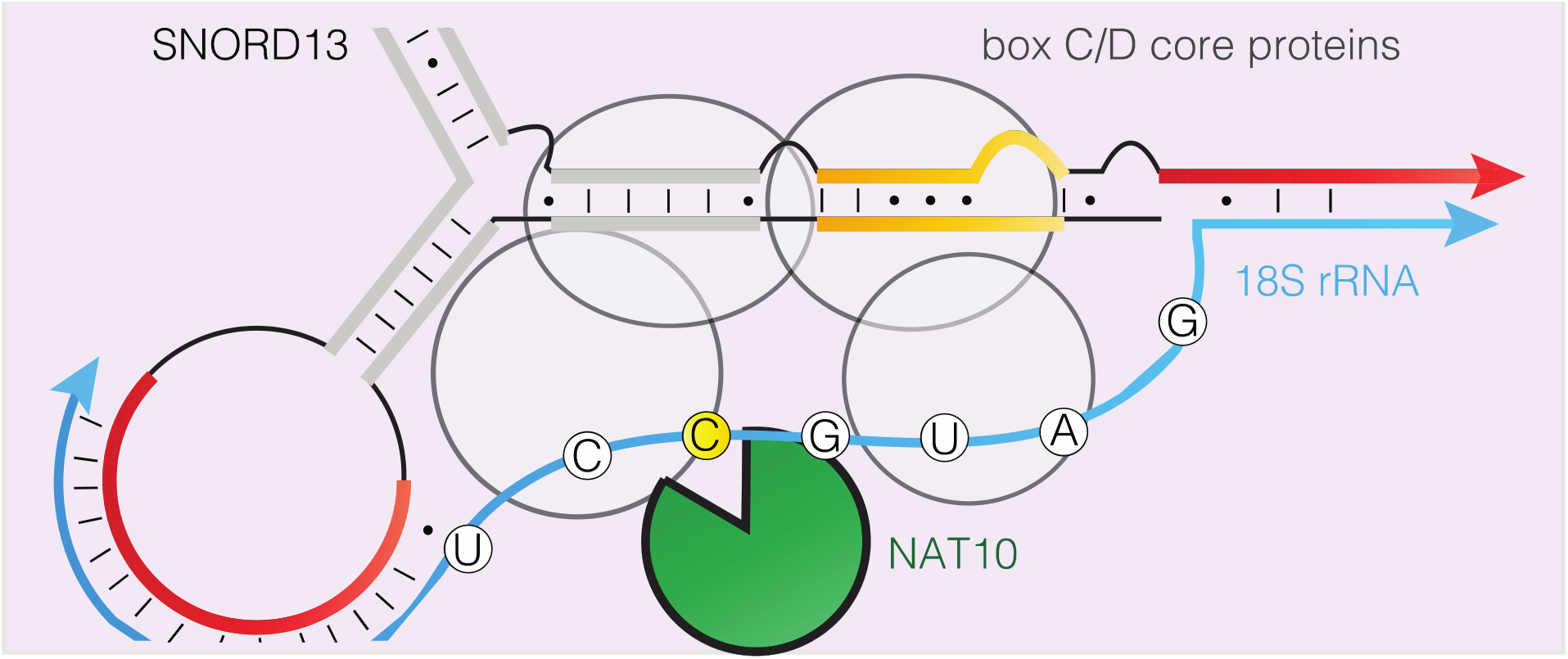

## Introduction

More than 100 modified RNA nucleotides are known across all domains of life. One of the most highly conserved of these is N4-acetylcytidine (ac^4^C).^*1*^ In humans, ac^4^C has been mapped to two dominant sites in serine and leucine tRNA, as well as two dominant sites in small subunit (SSU) 18S rRNA occurring at helix 34 and 45.^*2, 3*^ Acetylation of all of these RNA targets is catalyzed by NAT10, the only RNA acetyltransferase known in the human genome. NAT10’s selectivity towards tRNA or rRNA substrates is governed by the protein adapter THUMPD1 and the short nucleolar RNA (snoRNA) adapter SNORD13, respectively.^*3*^

Recently, we used genetic ablation to demonstrate that SNORD13 is required for acetylation of a single cytidine residue in helix 45 (h45) of small subunit 18S ribosomal RNA (SSU-ac^4^C1842; Figure 1a).^*4*^ In addition to its biological function, another question is how SNORD13 helps NAT10 recognize its targets in human cells. SNORD13 is a box C/D snoRNA, so termed for the presence of C (5’–RUGAUGA-3’) and D (5’–CUGA-3’) motifs in its primary sequence. However, several features of SNORD13 are atypical of this RNA class. First, while many box C/D snoRNAs play an important role in rRNA 2’-O-methylation by nucleating the assembly of short nucleolar ribonucleoprotein (snoRNP) complexes and guiding the 2’-O-methyltransferase Fibrillarin (FBL) to its target sites via antisense pairing, SNORD13 is the only human box C/D snoRNA known to be involved in cytidine acetylation.^*5*^ Second, SNORD13 is one of only three snoRNAs known to be independently transcribed by RNA Polymerase II (Pol II) in human cells.^*6*^ Third, SNORD13 contains two rRNA antisense elements, one located at the 5’ end and the other positioned more internally along the SNORD sequence (Figure 1b). It is also currently unclear to what extent the mechanisms box C/D snoRNAs use to guide cytidine acetylation are conserved across evolution. For example, the yeast SNORD13 homologue snr45 directly associates with the RNA acetyltransferase Kre33, and may belong to a larger C/D box snoRNP containing the FBL homologue Nop1.^*7*^ However, vertebrate SNORD13 shows little homology with snr45 (vide infra) and has a longer antisense region and shorter intervening structural loop. Another contrast with the yeast system comes from the fact that co-immunoprecipitation studies have not revealed a direct interaction of NAT10 and SNORD13 in human cells, despite many attempts.^*3, 8*^ Finally, homologues of the snoRNA that guides helix 34 cytidine acetylation in yeast (snr4) have yet to be identified in humans, and whether such guide mechanism exists remains unknown.

**Figure 1.**
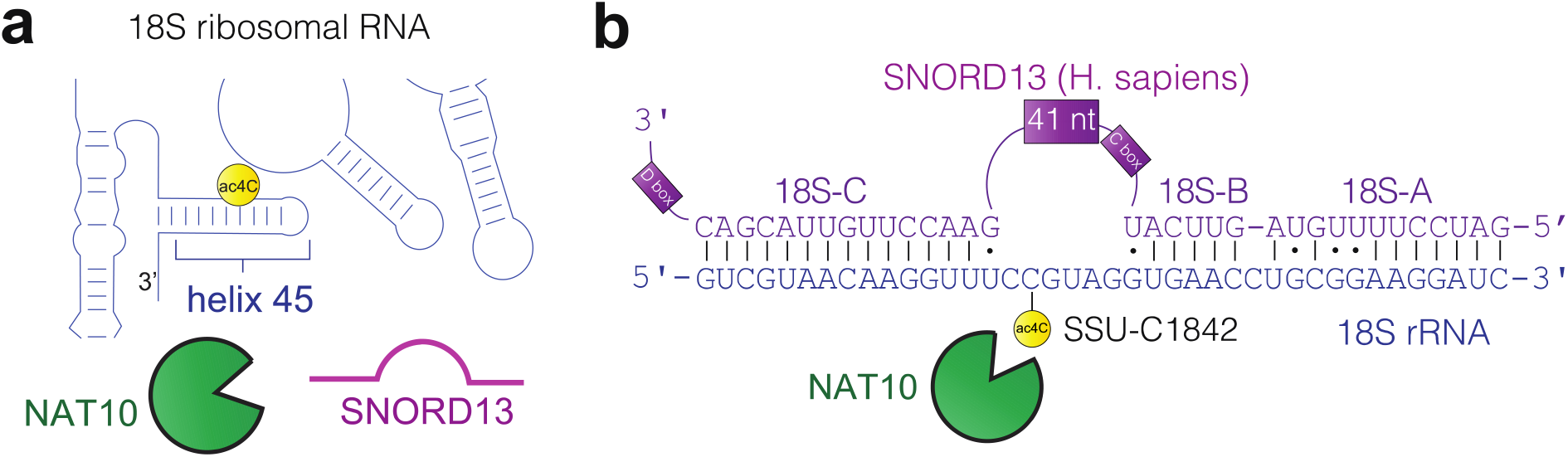
(a) NAT10 and SNORD13 are required for deposition of ac^4^C in helix 45 of 18S rRNA. (b) Schematic of antisense complementarity between human SNORD13 (purple) and 18S rRNA (blue). The site of cytidine acetylation (C1842) is specified.

A major challenge to understanding box C/D snoRNP function has been the difficulty of their in vitro reconstitution, which has heretofore been limited to archaeal enzymes. ^*9–11*^ In contrast, the study of eukaryotic systems has been enabled by the development of cellular reconstitution assays.^*12–15*^ In this strategy, mutants of an endogenous snoRNA are expressed ectopically in cells and tested for their ability to guide modification of a native (or mutated non-native) substrate. Applied to study the box C/D snoRNAs involved in ribose methylation, this approach has enabled the identification of motifs critical for snoRNA stability, rules for substrate targeting, and guidance of ribose methylation to novel rRNA sites in yeast as well as mammalian systems.^*13, 16, 17*^ To date, a single study has used this approach to study snoRNA-guided ac^4^C in yeast.^*7*^ Similar systems to study vertebrate cytidine acetylation have yet to be developed or applied.

Here we report two experimental approaches to probe SNORD13-dependent RNA acetyltransferase activity in human cells. Our first strategy hinges on the use of a SNORD13 knockout cell line in which SSU-ac^4^C1842 is lost.^*4*^ Ectopic expression of SNORD13 rescues h45 acetylation and can be quantitatively assessed using nucleotide resolution ac^4^C sequencing. Using this assay we evaluated several SNORD13 mutants, allowing us to identify structural and antisense elements critical for activity. To study SNORD13’s substrate specificity we developed a second approach. This assay uses a RNA Polymerase I (Pol I) transcribed minigene to express a nucleolar pre-rRNA fragment which is efficiently acetylated by endogenous NAT10/SNORD13. Systematic mutation of this substrate is used to validate a consensus sequence necessary for ac^4^C deposition and highlights the potential reprogrammability of this system. Our studies demonstrate the power of combining mutational analysis with nucleotide resolution ac^4^C sequencing, and provide the basis for understanding and exploiting RNA-guided RNA modification catalysts in biology, biotechnology, and disease.

## Results

### Conserved secondary structure of vertebrate SNORD13

Towards the goal of defining functional elements in SNORD13, we first analyzed this snoRNA for evidence of conserved secondary structures (Figure 2a). All SNORD13 genes contain canonical C- and D-boxes (yellow) as well as antisense segments with complementarity to 18S rRNA (red); however, little sequence homology is evident outside these regions. Of note, the regions of complementarity of human SNORD13 contain bulges and unpaired bases and architecturally are not identical to SNORD13-like RNAs found in *A. thaliana* (snoR105, snoR108 and snoR146)^*18*^ and *S. cerevisiae* (snr45)^*7*^. For clarity, we refer to SNORD13’s antisense elements as 18S-A and 18S-B (located at the 5’-end) and 18S-C (positioned more internally; Figure 1b, 2a).

**Figure 2.**
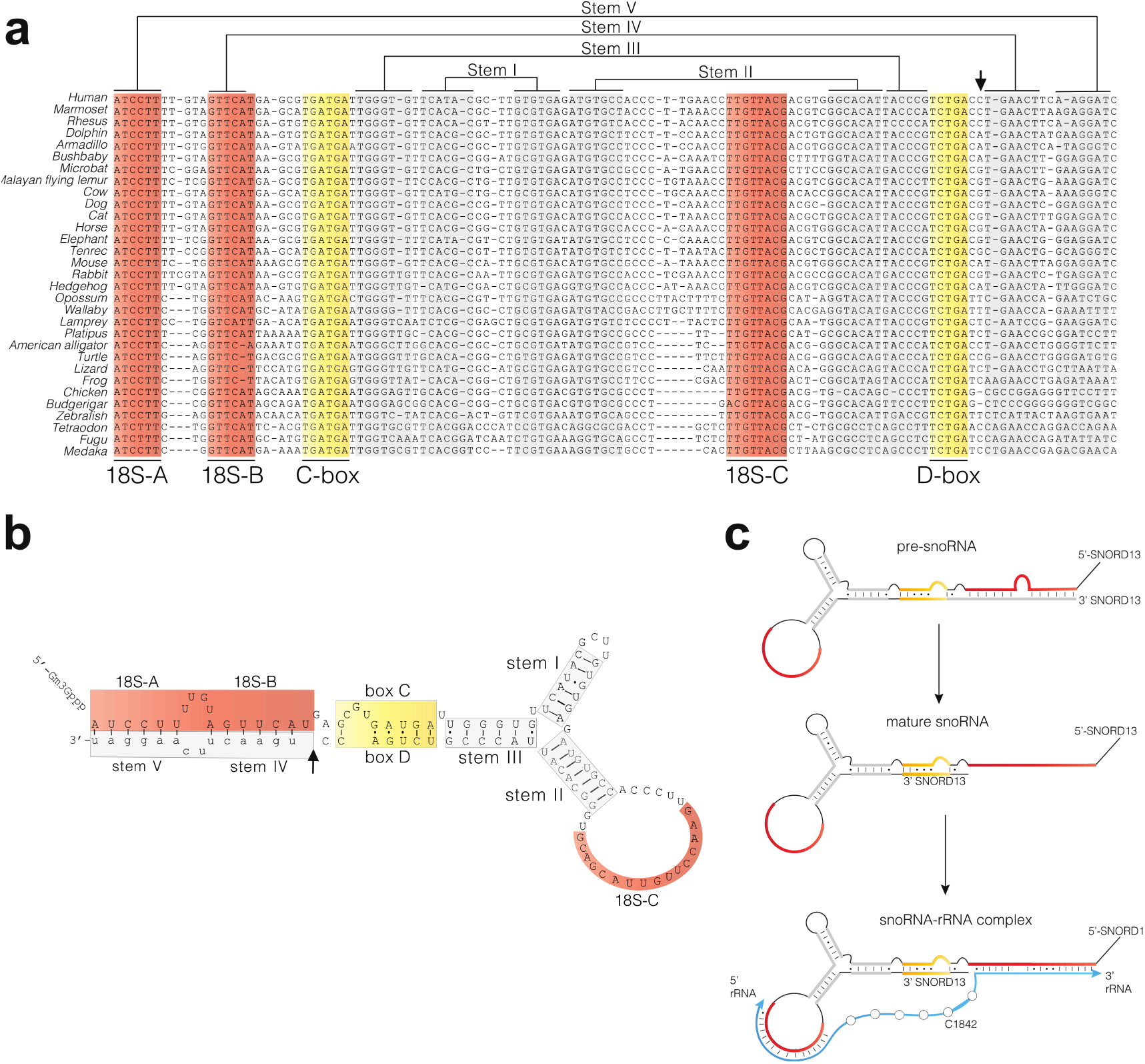
(a) Sequence alignment of vertebrate SNORD13 homologues. Antisense regions 18S-A, 18S-B, and 18S-C are highlighted in red, C and D-box motifs involved in k-turn formation in yellow, and putative stem regions in grey. The small verticle black arrow indicates the 3’ end of SNORD13 as experimentally determined by RACE analysis. (b) Proposed SNORD13 secondary structure with colors coded as above. Bases written in uppercase are part of stable, fully processed SNORD13 while those written in lowercase correspond to 3’-extension of the transient pre-SNORD13 intermediate. (c) Proposed maturation of pre-SNORD13, containing a 3’-extension which masks its 5’-antisense region, to mature SNORD13 capable of interacting with 18S rRNA.

Taking into account formation of the kink-turn motif, previously found to be crucial for box C/D snoRNP assembly,^*19*^ *in silico* structural analysis predicted three short, conserved stems in SNORD13 which we designate as stems I, II and III, respectively (Figure 2b). We further observed that the 18S-A and 18S-B antisense elements have the potential to base-pair with an ~15 nucleotide region downstream of the 3’-end of mature SNORD13, defining two additional helices which we refer to as stem IV and stem V, respectively (Figure 2b). With the exception of Lamprey these latter two stems appear less conserved in non-eutherian species. The formation of stem IV and stem V requires that transcription of SNORD13 gene goes beyond its mature 3’ - end, which has been previously observed.^*20–22*^ These external stems may be important to formation of the kink-turn motif by bringing together the C- and D-boxes,^*23*^ or serve as signal that delineates the mature 3’-end. Of note, a process analogous to the latter has been recently found to play a role in regulating expression of SNORD118.^*24, 25*^ Overall, our analyses suggest fully processed SNORD13 folds into three evolutionarily conserved stems which scaffold three distinct 18S rRNA antisense elements, two of which are unmasked from a 3’-extended precursor to enable facile base-pairing with rRNA (Figure 2c).

### Structural elements in stem I, III, and IV dictate SNORD13 expression

Secondary structure plays a critical role in snoRNA processing, protein interactions, and cellular accumulation.^*26, 27*^ To begin to build sequence-function relationships for SNORD13, we first examined how mutating the secondary structural elements predicted above affected its stability in cells (Figure 3a). To achieve this objective, we transfected a 1200 nt-long genomic human DNA fragment predicted to contain all *cis*-acting elements required for transcription and 3’-end processing of SNORD13. Our experimental strategy used 11 human SNORD13 constructs comprising four groups: i) the wild type SNORD13 sequence, ii) two mutants designed to disrupt the C and D boxes, iii) four mutants designed to disrupt stems I, II, III, and IV, and iv) four mutants designed to rescue complementarity in stem-disrupting mutations (Figure 3b). In order to distinguish ectopically expressed human SNORD13 from its endogenously expressed mouse counterpart, each construct was individually transfected into mouse L929 cells. SNORD13 expression was then assessed 48 hours post-transfection by ribonuclease (RNAse A/T1) protection assay using ^*32*^P-labelled antisense riboprobes specific for each mutated version of SNORD13 (Figure 3a). Accordingly, protected RNA fragments in the top panel of Figure 3c, which are only detected in transfected samples, represent full length transfected version of SNORD13. Those in the bottom panel, which are detected in both transfected and non-transfected samples, originate from partial protection of endogenous background RNAs (including mouse SNORD13, due to sequence divergence between human and mouse), providing a rough measure of loading.

**Figure 3.**
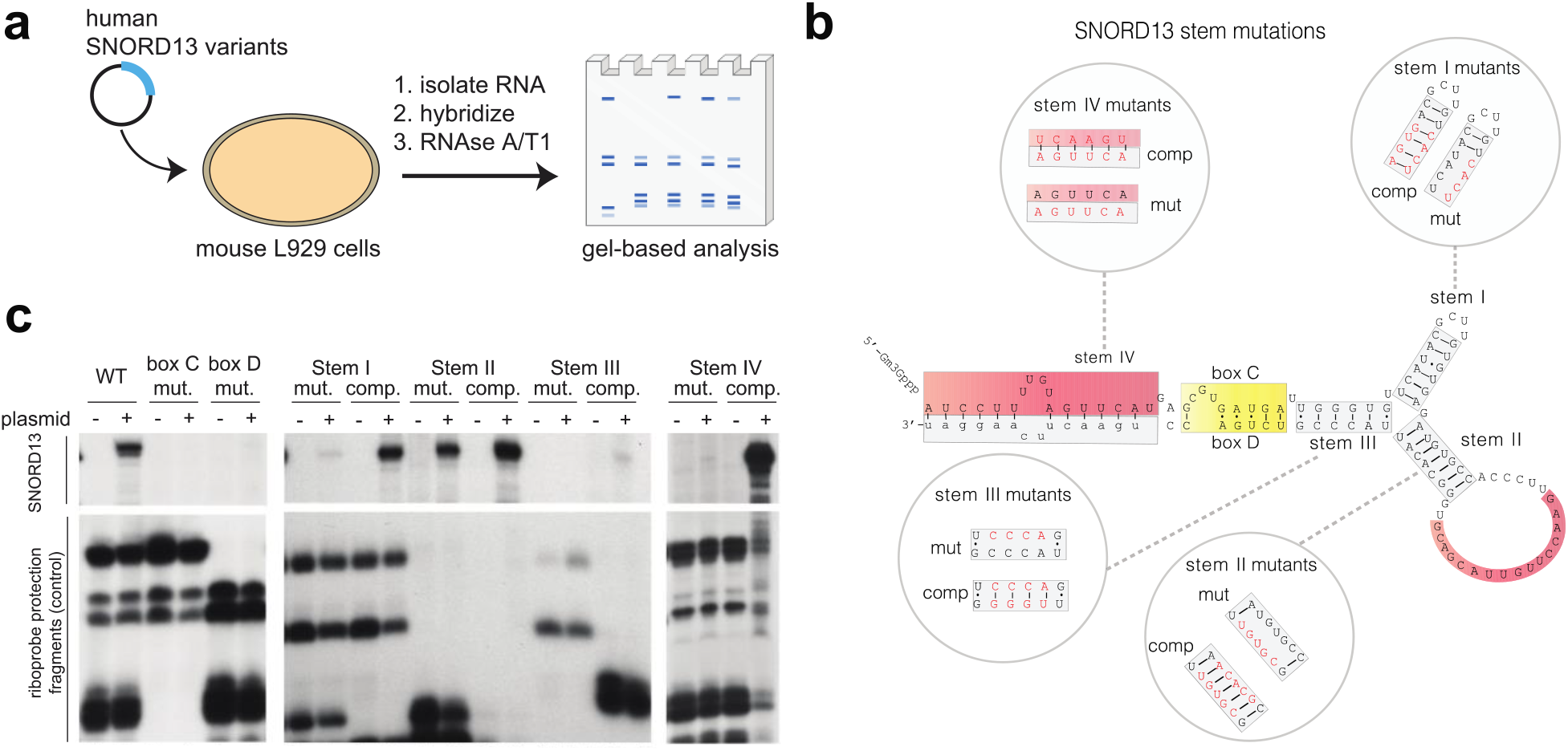
(a) Schematic for analysis of SNORD13 stability by RNAse A/T1 assay. (b) Sequence of human SNORD13s analyzed. In addition to wild-type (“WT,” black) sequence, disruptive mutations (“mut,” red), and rescue mutations (“comp,” red) were explored for stems I-IV. C and D box mutations were also analyzed (sequences provided in Supplementary Information). (c) Results of RNAse A/T1 stability assay. Top box indicates protected RNA fragment corresponding to full length, transfected human versions of SNORD13. Bottom box indicates protected RNA fragments corresponding to endogenously-expressed mouse SNORD13 background.

Mutation of either the C or D box abolished detection of SNORD13 (Figure 3c), consistent with previous analyses showing the C/D box is critical for snoRNP formation and snoRNA accumulation.^*26, 27*^ Mutations causing disruption of stems I, III and IV also led to a complete loss of observed SNORD13 expression. In the cases of stems I and IV, SNORD13 levels could be restored by introducing complementary mutations that rescue the anticipated helical structures (Figure 3b-c). However, stem III showed minimal rescue by this approach, suggesting a critical role for its sequence in SNORD13 folding or function (Figure 3c). Previous studies have proposed the formation of a pseudo-knot between nucleotides in stem III and 18S-C in the yeast SNORD13 homologue may be required to bring the antisense regions into close proximity,^*7*^ and it is possible complementary mutations fail to rescue this structure in SNORD13. Mutations that disrupt stem II and flanking the 18S-C antisense element were well-tolerated and did not appear to affect SNORD13 abundance (Figure 3c). This may imply stem II does not form or, alternatively, that it is dispensable for the processing and stability of SNORD13. These studies provide experimental evidence for the existence of a C/D box and RNA secondary structures in human SNORD13 that are critical to its cellular accumulation. Furthermore, we observe that functional features of SNORDI3’s predicted structure, such as the occurrence of antisense elements in single-stranded regions, are shared with yeast. This implies that despite low sequence homology, the structure of human SNORD13 may resemble that of yeast snr45.

### Rescue of cytidine acetylation depends on SNORD13 structure and base-pairing

In an accompanying study,^*4*^ we determined that inactivation of SNORD13 in the human chronic myelogenous leukemia-derived HAP1 cell line causes loss of SSU-ac^4^C1842 in 18S rRNA. To understand the molecular determinants of this process, we next evaluated the ability of several ectopically-expressed SNORD13 mutants to rescue SSU-ac^4^C1842 in this cell line (Figure 4a). Briefly, SNORD13-deficient HAP1 cells were co-transfected with two plasmids expressing SNORD13 and eGFP. After 48 hours, flow cytometry was used to isolate cells expressing eGFP (presumably also enriched in SNORD13 expression), and RNA from these cells was analyzed by nucleotide resolution ac^4^C sequencing. This method uses a chemical reaction that modifies the structure of ac^4^C residues, leading to misincorporations (C-to-U) when this RNA is reverse transcribed.^*2, 8*^ PCR amplification and sequencing are then used to provide a quantitative readout of ac^4^C’s presence in RNA at a position of interest.

**Figure 4.**
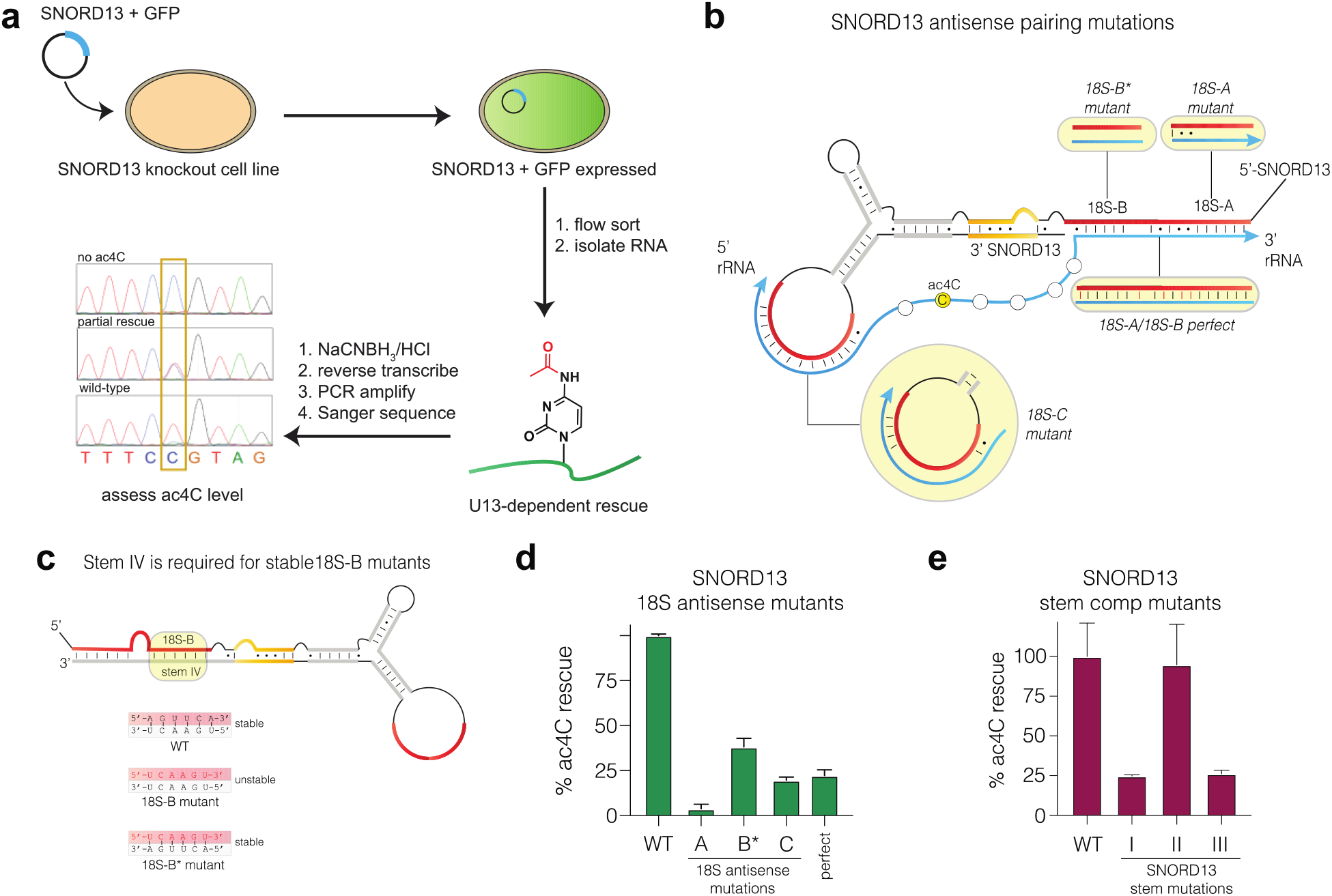
(a) Schematic for analysis of SNORD13 function using quantitative ac^4^C sequencing. (b) Sequence of human SNORD13s analyzed. In addition to wild-type ("WT") sequence, disruptive mutations (18S-A, 18S-B*, 18S-C mutants), and a mutant with increased complementarity ("18S-A/18S-B perfect comp") were explored for rescue of ac^4^C in SNORD13 KO cells. Sequences and verification of expression are provided in Figure S1. Note the finding that the 18S-A mutant is expressed when its stem V is disrupted, implies stem V is not strictly required for SNORD13 synthesis. (c) Stem IV is required for accumulation of mutant SNORD13s. Mutation of SNORD13 in mutant 18S-B disrupts stem IV. Re-introduction of complementarity in construct 18-B* (bottom) allows accumulation and testing of function. Expression of 18S-B* is verified by RNAse A/T1 mapping in Figure S1c. (d) Rescue of SSU-ac^4^C1842 by SNORD13 antisense mutants. Mutants rescue values are normalized relative to the WT SNORD13, which was set to equal 100%. Background misincorporation rates in SNORD13 KO cells were 0-4%. Exemplary sequencing traces are provided in Figure S1d. (e) Rescue of SSU-ac^4^C1842 by SNORD13 stem comp mutants. Structures of comp mutants are provided in Figure 3b.

To explore whether SNORD13’s antisense guides are essential for acetylation, we compared ac^4^C rescue by wild-type SNORD13 to three mutants in which 18S-A, 18S-B, and 18S-C were disrupted (18S-A, 18S-B, and 18S-C mutants; Figure 4b, Figure S1a). Since 18S-B’s antisense element is part of stem IV, whose disruption destabilizes SNORD13 (Figure 3c), we used a second-generation mutant of 18S-B (termed 18S-B*) which restores formation of stem IV (Figure 4c, Figure S1b-c). As expected, rescue of SSU-ac^4^C1842 by ectopically-expressed wild-type SNORD13 was readily observed using ac^4^C sequencing (Figure 4d, Figure S1d). Disruption of antisense complementarity diminished rescue by 60-92%, with the deleterious effect of mutations following the order 18S-A >> 18S-C > 18S-B (Figure 4d, Figure S1d). The finding that 18S-A, which covers nt 1-6 of SNORD13 and base-pairs to nts 1854-1859 of 18S rRNA, had the most profound effect on SNORD13 activity is consistent with observations in yeast^*7*^ but contrasts with our recent analysis of SNORD13 homologues in Drosophila,^*4*^ where 5’ antisense elements do not appear to be found. To more thoroughly probe the impact of antisense complementarity on ac^4^C rescue, we designed a SNORD13 mutant which we "perfected" by correcting five mismatched or unpaired bases found in the 18S-A and B regions (Figure 4b; ‘18S-A/18S-B perfect comp’). However, compared to wild-type SNORD13, this mutant also exhibited reduced rescue of SSU-ac^4^C1842. Previous studies of C/D box snoRNAs that target FBL have found methylation levels do not necessarily correlate with the stability of the substrate-guide duplex or snoRNA levels, and may be subtly tuned by a complex interaction network depending on the sequence of its RNA substrate.^*9, 28*^ Consistent with this notion, the 18S-C mutant appears more weakly expressed than 18S-A (Figure S1c), yet displays more effective rescue of RNA acetylation. Finally we applied this assay to examine the effect of stem I, II, and III on rescue, exclusively focusing on mutations that maintain stem complementarity (Figure 3b; ‘comp’). Replacement of stems I and III with stems of equivalent length but altered sequence did not rescue ac^4^C, suggesting a functional role for these elements (Figure 4e), with the caveat that stem III comp may be poorly expressed (Figure 3c). In contrast, expressing a mutant SNORD13 with altered sequences in stem II appears well tolerated (Figure 4e). Thus, the sequence forming stem II appears dispensable for both U13 stability and NAT10 substrate targeting. Overall, our studies reveal features of SNORD13 critical to guiding ac^4^C deposition and highlight loss of substrate-guide complementarity at the 5’ end (nts 1-6) as a distinctly disruptive perturbation.

### Sequence requirements of SNORD13 substrates

The presence of multiple copies of genes encoding ribosomal RNA presents a challenge to effective mutagenesis and hinders analysis of the substrate specificity of snoRNA-guided RNA modification in human cells. Beyond this technical barrier, prior attempts to mutagenize SSU-ac^4^C substrates in yeast found every mutation introduced in helix 45 of 18S rRNA to be lethal.^*7*^ Seeking to circumvent these limitations, we were inspired by the utility of ribosomal minigenes expressing the 3’ terminus of 18S rRNA fused to internal transcribed spacer 1 (referred to here as h45-ITS1) in investigating the role of SNORD13 in pre-rRNA processing.^*17, 29, 30*^ The expected recognition of the h45-ITS1 minigene by the pre-rRNA processing machinery led us to hypothesize it may also be able to interact with endogenous NAT10/SNORD13, enabling its facile mutagenesis for substrate specificity studies. To test this we expressed an h45-ITS1 minigene construct in HEK-293T cells, isolated RNA, and analyzed it using the aforementioned ac^4^C sequencing workflow (Figure 5a). Importantly, these experiments used primers that specifically amplify minigene h45-ITS1, rather than the endogenous 18S pre-RNA (Figure S2a-b). Ribosomal minigenes expressed from a RNA Polymerase I (Pol I)^*31*^ but not an RNA Polymerase II (Pol II) promoter were efficiently acetylated, consistent with the predominant nucleolar localization of NAT10 and SNORD13 (Figure S2c). Modification occurred at the expected position corresponding to SSU-C1842 of 18S rRNA. Of note, detection of ac^4^C in this ectopically-expressed pre-rRNA fragment requires the use of the nucleotide resolution sequencing assays, which enables PCR amplification of the ac^4^C signal and represents an advance over previous analytical methods.

**Figure 5.**
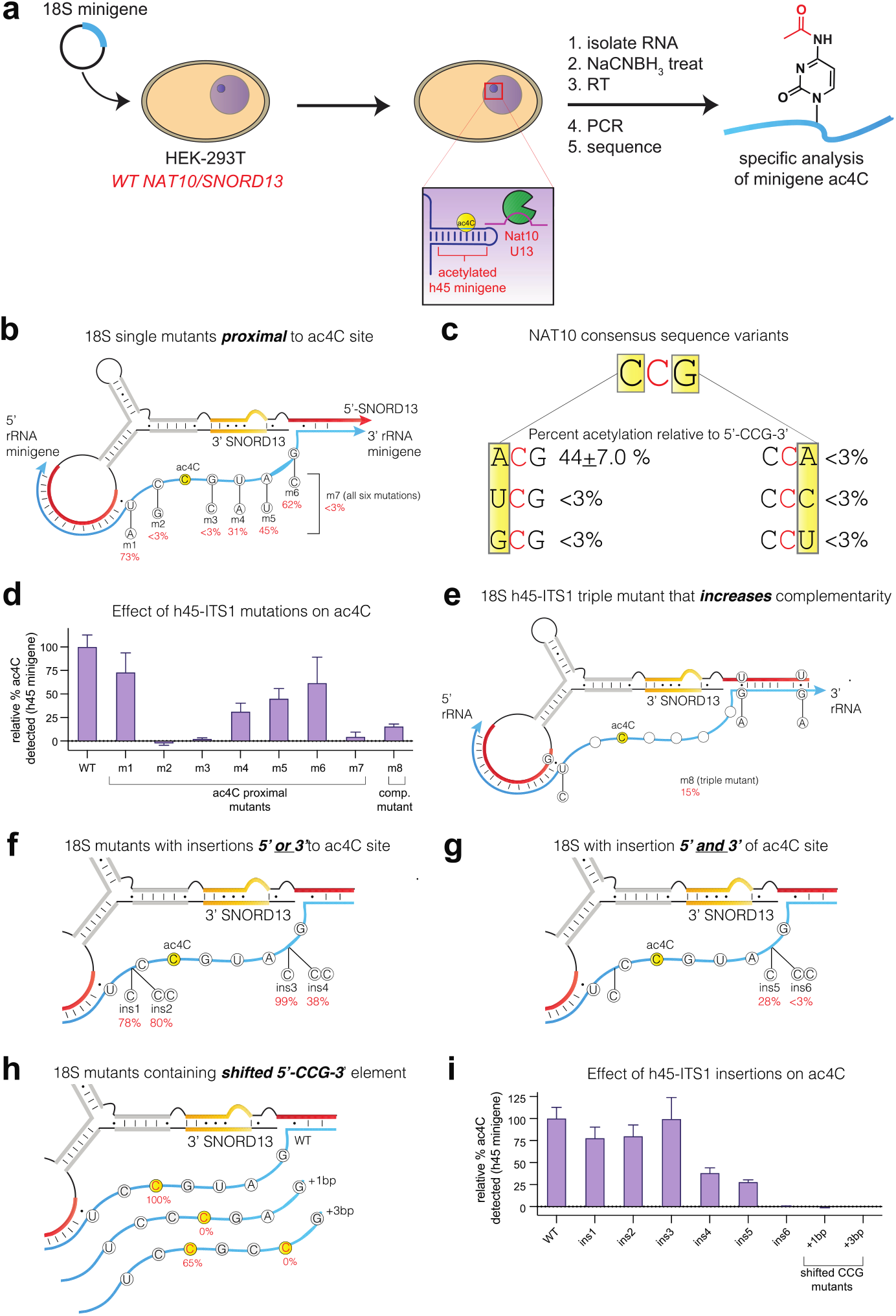
(a) Schematic for analysis of endogenous SNORD13 substrate specificity using a Pol I-transcribed pre-rRNA h45-ITS1 minigene substrate and quantitative ac^4^C sequencing. (b) Structure of h45-ITS1 substrates with mutations lying proximal to natural ac^4^C site. Values in red represent ac^4^C-dependent misincorporation rates normalized relative to the WT h45-ITS1 sequence, which was set to equal 100%. (c) Summary of percent misincorporation observed upon mutation of 5’-CCG-3’ consensus sequence in h45-ITS1 substrates. (d) Bar graph of mutants specified in Figure 5b and 5e. (e) Structure of ‘triple mutant’ h45-ITS1 substrate engineered to have increased complimentarity to SNORD13. (f-g). Structure of h45-ITS1 substrates with bases inserted 5’ and/or 3’ relative to the 5’-CCG-3’ consensus sequence. (h) Structure of h45-ITS1 substrates in which the 5’-CCG-3’ consensus sequence is shifted 1 bp (+1bp) or 3 bp (+3bp). i) Bar graph of mutants specified in Figure 5f-h. Exemplary sequencing traces provided in Figure S3 and S5b.

To begin to understand the substrate specificity of SNORD13-dependent RNA acetylation we analyzed h45-ITS1 constructs in which individual residues flanking SSU-C1842 were mutated but SNORD13 complementarity was preserved (Figure 5b-c). Previous studies have shown eukaryotic NAT10 demonstrates a strong preference for modification of 5’-CCG-3’ sequences.^*2, 32, 33*^ Whether SNORD13-dependent acetylation is able to tolerate mutation of this consensus remains unknown. Maintaining the central acetylation site and focusing on the 5’ and 3’ residues (Figure 5c), we unexpectedly observed significant acetylation (~44% misincorporation) of a non-canonical 5’-ACG-3’ sequence. Re-analysis of a previously reported degenerate substrate library furnished further evidence that human NAT10 can modify 5’-ACG-3’ as well as 5’-UCG-3’ sequences when massively overexpressed in HEK-293T cells (Figure S4), albeit with greatly reduced efficiency relative to 5’-CCG-3’. In contrast, further changes-including all substitutions of the 3’ guanosine-reduced modification of the h45-ITS1 pre-rRNA to negligible levels (<3% misincorporation). Evaluating constructs with mutations lying outside of the consensus, we found that 5’ mutations (m1) were seemingly more well-tolerated than those lying 3’ (m4/m5). Simultaneous mutation of all six residues completely eliminated modification (m7), as expected (Figure 5b-d). Mutations in h45 designed to improve snoRNA base-pairing (m8) paradoxically lessened modification of the h45-ITS1 substrate (Figure 5d-e). This is consistent with our analysis of SNORD13 above (Figure 4b-d; ‘18S-A/18S-B perfect comp’), and further emphasizes that ac^4^C deposition does not strictly correlate with the apparent strength of guide-substrate complementarity.

Many snoRNA-dependent RNA modification systems exhibit strict spatial requirements for catalysis. For example, box C/D snoRNAs direct ribose methylation to the nucleotide base-paired to the fifth residue upstream of the D or D’ boxes, and insertion of an intervening base can shift what nucleotide is methylated.^*13, 16*^ H/ACA box snoRNAs contain two antisense elements that form a three-way junction with substrates, leaving the targeted uridine and a flanking residue unpaired and presented to the pseudouridine synthase in the center of a pseudouridinylation pocket which occurs a defined distance (14-16 nt) from the H box or ACA box.^*34–36*^ These strict spatial requirements reflect the precision with which guide snoRNAs must properly position their substrates in the active site of modification enzymes to enable catalysis. To probe whether similar requirements exist for SNORD13-dependent RNA acetylation, we inserted nucleotides in the putatively unpaired ‘acetylation pocket’ of the h45-ITS1 pre-RNA substrate (Figure 5f-i). Addition of one or two residues 5’ to the modification site (ins1, ins2; Figure 5f) had little affect on ac^4^C deposition (Figure 5i). Inserting a single nucleotide 3’ to the modification site (ins3) was also tolerated. However, inserting two nucleotides 3’ to the modification site of h45 (ins4) led to a >50% loss of acetylation, and when combined with an additional upstream cytidine (ins6) abolished modification entirely (Figure 5f-i). The finding that the ins4 construct contains an additional 5’-CCG-3’ that is not modified implies the relative location of the 5’-CCG-3’ consensus sequence matters. To test this we attempted to shift the position of the 5’-CCG-3’ consensus without altering the overall length of the loops (+1bp and +3bp mutants; Figure 5h). These mutant substrates were not modified at the novel 5’-CCG-3’ sequence; however, in the +3bp mutant, a degree of modification (65%) was observed at the site corresponding to SSU-C1842 (Figure 5h, Figure S5). Interestingly, our studies found disruption of 18S-B complementarity enables a degree of SNORD13-dependent ac^4^C deposition (Figure 4d). This would seemingly preclude a rule in which a defined number of intervening bases must lie between the antisense-paired rRNA substrate and nucleotide to be modified, although additional possibilities exist. Overall, our studies indicate unusual plasticity in the NAT10 active site which accommodates diverse presentation of 5’-HCG-3’ (H = A/C/U) pre-rRNA substrates at the endogenous C1842 position.

Previous studies have exploited designer snoRNAs to direct modifications to novel RNA targets.^*16*^ This has been used as an alternative to mutagenesis to explore ribosome function,^*37, 38*^ and to influence modification-sensitive processes such as nonsense suppression.^*39*^ As a simple proof-of-concept, we tested whether a mutant snoRNA could increase acetylation of an h45-ITS1 substrate that is poorly recognized by endogenous SNORD13 (Figure 6a). Mutation of h45-ITS1 to disrupt its interaction with 18S-A (nts 1-6) largely abrogated the ability of endogenous SNORD13 to guide its modification, leading to an ~90% decrease in ac^4^C (Figure 6b). Ectopic expression of a SNORD13 containing an 18S-A region designed to base-pair with the mutant substrate partially rescued its modification. Although additional applications of this system lie beyond the scope of our current study, this experiment demonstrates the potential of our assays to explore how cytidine acetylation may be directed to non-native substrates using synthetic SNORD13 analogues.

**Figure 6.**
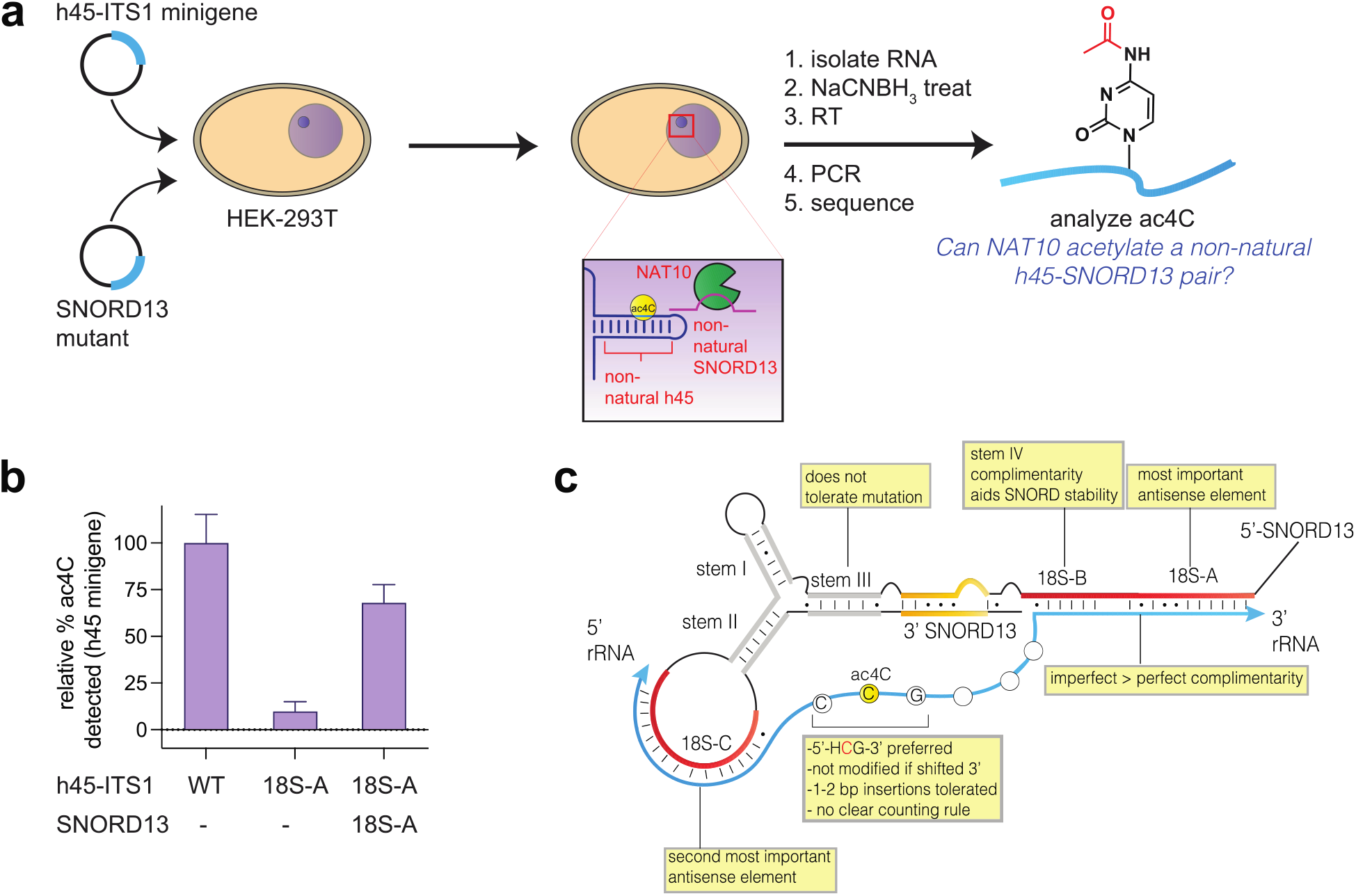
(a) Schematic for analysis of orthogonal SNORD13-substrate pairs. Exemplary sequencing traces are provided in Figure S6. (b) Bar graph of ac^4^C levels at site corresponding to SSU-1842 in orthogonal SNORD13-substrate pairs. Values represent ac^4^C-dependent misincorporation rates normalized relative to the WT h45-ITS1 and WT SNORD13 pair, which was set to equal 100%. Sequences of SNORD13 18S-A mutant and h45-ITS1 18S-A mutant are provided Supplementary Information. (c) Overview of SNORD13 structure-function relationships probed in this study.

## Discussion

Here we have reported the development of parallel experimental strategies to study SNORD13-dependent RNA acetylation in human cells. We demonstrate that SNORD13 knockout cells provide an optimal genetic background to examine rescue of SSU-ac^4^C1842 by ectopically-expressed snoRNAs, while h45-ITS1 ribosomal minigenes can be used to profile the substrate selectivity of endogenous SNORD13 (Figure 6c). Quantitative analysis of each of these systems is enabled by a signal amplified ac^4^C sequencing method. Applying these assays highlighted a critical role for stem I, stem III, and the 18S-A antisense regions in SNORD13 activity and validated the preference for substrate acetylation to occur in a 5’-CCG-3’ consensus sequence, with 5’-ACG-3’ also tolerated. These systems also allowed us to probe spatial constraints in the putative pocket formed by the snoRNP responsible for acetylation, which appears to tolerate insertion of unpaired nucleotides upstream, but not downstream, of the SSU-C1842 modification site.

A technical limitation of our studies is the precision of Sanger sequencing-based ac^4^C detection, which as used here is most conducive to relative comparisons of SNORD13 activity (detecting differences of ~25%) rather than absolute quantitative analysis. In the future this may be improved by integrating our methods with next-generation sequencing workflows, ^*2, 40*^ allowing finer scale differences to be observed. Future studies may also benefit from the use of alternative methods to knockdown SNORD13, for example using modified antisense oligonucleotides (ASOs),^*31*^ which could enable the functionality of SNORD13 analogues to be tested in new cell lines and model organisms not amenable to genetic manipulation.

The composition and mechanism of the snoRNP which catalyzes SSU-ac^4^C1842 remain unknown. While both NAT10 and SNORD13 are clearly required for h45 acetylation, we and others^*3*^ have not yet found evidence that the acetyltransferase and snoRNA to directly interact. Indeed, the most well-characterized protein binding partner of SNORD13 is FBL,^*6, 41*^ suggesting its likely integration into a C/D snoRNP comprised of 15.5K, NOP56, and NOP58.^*19, 42*^ Yeast NAT10 has been proposed to co-occupy snr45 together with this organism’s C/D snoRNP protein homologues,^*7*^ and interactions between NAT10, FBL, and NOP56 identified in recent proteome-wide studies raise the possibility of a similar scenario in human cells.^*43*^ One model for this to occur would be if a core snoRNP comprising SNORD13 bound to 18S rRNA, recruited NAT10 via a transient protein-protein interaction, and presented C1842 to its active site for acetylation. The in vitro characterization of NAT10-containing snoRNPs may benefit from the development of simplified archaeal systems, as has been useful in studies of snoRNA-dependent methylation and pseudouridinylation.^*2, 11, 35*^ For the moment, the assays reported here should provide a powerful tool with which to determine the contribution of different snoRNP components to SSU-C1842 acetylation in human cells.

An additional motivation for studying snoRNAs lies in their potential to be engineered to direct RNA modifications to novel targets.^*16, 37*^ As an initial step in this direction, we demonstrated that a SNORD13 mutant can be used to drive acetylation of an ectopic pre-rRNA substrate that is not efficiently modified in endogenous cells. It is important to specify that this represents an extremely simple model system, and the extent of SNORD13’s reprogrammability, as well as whether analogues may direct NAT10 to novel substrates, remains unknown. Considering our collective observations, the following features of SNORD13 appear necessary for ac^4^C deposition: i) antisense complementarity, with apparent importance following 18S-A > 18S-C, 18S-B, ii) invariable stem I and III sequences, iii) a stem IV engineered to complement 18S-B for stability, iv) a 5’-MCG-3’ substrate sequence (M=A or C), and v) a non-hybridized substrate loop which tolerates flexibility but not shifting of the cytidine modification site within it (Figure 6c). The tolerance for a 5’ C→A mutation in the CCG consensus is in line with our recent report of acetylation in a 5’-UCG-3’ sequence in the model organism *P. polycephalum*.^*4*^ In contrast, substrates harboring mutations 3’ to the acetylation site (e.g. 5’-CCC-3’) were not modified. This suggests human NAT10 strongly prefers modification of 5’-HCG-3’ sequences, a consideration that should be taken into account when evaluating novel substrates.^*2, 44*^ It has been previously noted that SNORD13 resembles a H/ACA snoRNA in terms of its bipartite substrate complementarity.^*7, 45*^ Our mutational analyses of the antisense regions of SNORD13 - as well as our recent discovery of an atypical *D. melanogaster* SNORD13 homologue^*4*^ - suggest that unlike H/ACA snoRNAs, a single antisense strand can suffice for ac^4^C deposition while the second may increase efficiency. Additional studies will be required to characterize this model and establish the scope of substrates NAT10 may be directed to via mutated SNORD13 analogues. Such research will be greatly aided by high-throughput methods to characterize RNA-RNA interactions^*46*^ as well as the transcriptome-wide distribution of ac^4^C.^*2*^ Such studies are in progress and will be reported in due course.

## Acknowledgements

This work was supported by the Intramural Research Program of the NIH, National Cancer Institute, Center for Cancer Research (ZIA-BC011488 to J.L.M.), the Agence Nationale de la Recherche (ANR-18-CE12-0008-01 to J.C.), and the European Research Council (ERC) under the European Union’s Horizon 2020 research and innovation programme (grant agreement No. 714023 to S.S.).

## Supplementary Information

Supplementary figures, including materials and methods and sequences for constructs used in reconstitution experiments, are available online.

## References

[1] Dunin-Horkawicz, S., Czerwoniec, A., Gajda, M. J., Feder, M., Grosjean, H., and Bujnicki, J. M. (2006) MODOMICS: a database of RNA modification pathways, Nucleic Acids Res 34, D145–149.

[2] Sas-Chen, A., Thomas, J. M., Matzov, D., Taoka, M., Nance, K. D., Nir, R., Bryson, K. M., Shachar, R., Liman, G. L. S., Burkhart, B. W., Gamage, S. T., Nobe, Y., Briney, C. A., Levy, M. J., Fuchs, R. T., Robb, G. B., Hartmann, J., Sharma, S., Lin, Q., Florens, L., Washburn, M. P., Isobe, T., Santangelo, T. J., Shalev-Benami, M., Meier, J. L., and Schwartz, S. (2020) Dynamic RNA acetylation revealed by quantitative cross-evolutionary mapping, Nature 583, 638–643.

[3] Sharma, S., Langhendries, J. L., Watzinger, P., Kotter, P., Entian, K. D., and Lafontaine, D. L. (2015) Yeast Kre33 and human NAT10 are conserved 18S rRNA cytosine acetyltransferases that modify tRNAs assisted by the adaptor Tan1/THUMPD1, Nucleic Acids Res 43, 2242–2258.

[4] Bortolin-Cavaille, M. L., Aurélie, Q., Gamage, S. T., Thomas, J. M., Sas-Chen, A., Sharma, S., Plisson-Chastang, C., Vandel, L., Blader, P., Lafontaine, D., Schwartz, S., Meier, J. L., and Cavaille, J. (2022) Probing small ribosomal subunit RNA helix 45 acetylation across eukaryotic evolution, bioRxiv.

[5] Bratkovic, T., Bozic, J., and Rogelj, B. (2020) Functional diversity of small nucleolar RNAs, Nucleic Acids Res 48, 1627–1651.

[6] Tyc, K., and Steitz, J. A. (1989) U3, U8 and U13 comprise a new class of mammalian snRNPs localized in the cell nucleolus, EMBO J 8, 3113–3119.

[7] Sharma, S., Yang, J., van Nues, R., Watzinger, P., Kotter, P., Lafontaine, D. L. J., Granneman, S., and Entian, K. D. (2017) Specialized box C/D snoRNPs act as antisense guides to target RNA base acetylation, PLoS Genet 13, e1006804.

[8] Thomas, J. M., Briney, C. A., Nance, K. D., Lopez, J. E., Thorpe, A. L., Fox, S. D., Bortolin-Cavaille, M. L., Sas-Chen, A., Arango, D., Oberdoerffer, S., Cavaille, J., Andresson, T., and Meier, J. L. (2018) A Chemical Signature for Cytidine Acetylation in RNA, J Am Chem Soc 140, 12667–12670.

[9] Graziadei, A., Gabel, F., Kirkpatrick, J., and Carlomagno, T. (2020) The guide sRNA sequence determines the activity level of box C/D RNPs, Elife 9.

[10] Omer, A. D., Ziesche, S., Ebhardt, H., and Dennis, P. P. (2002) In vitro reconstitution and activity of a C/D box methylation guide ribonucleoprotein complex, Proc Natl Acad Sci U S A 99, 5289–5294.

[11] Tran, E. J., Zhang, X., and Maxwell, E. S. (2003) Efficient RNA 2’-O-methylation requires juxtaposed and symmetrically assembled archaeal box C/D and C’/D’ RNPs, EMBO J 22, 3930–3940.

[12] Deryusheva, S., and Gall, J. G. (2013) Novel small Cajal-body-specific RNAs identified in Drosophila: probing guide RNA function, RNA 19, 1802–1814.

[13] Kiss-Laszlo, Z., Henry, Y., Bachellerie, J. P., Caizergues-Ferrer, M., and Kiss, T. (1996) Site-specific ribose methylation of preribosomal RNA: a novel function for small nucleolar RNAs, Cell 85, 1077–1088.

[14] Liang, W. Q., and Fournier, M. J. (1997) Synthesis of functional eukaryotic ribosomal RNAs in trans: development of a novel in vivo rDNA system for dissecting ribosome biogenesis, Proc Natl Acad Sci U S A 94, 2864–2868.

[15] Liang, X. H., Liu, Q., and Fournier, M. J. (2009) Loss of rRNA modifications in the decoding center of the ribosome impairs translation and strongly delays pre-rRNA processing, RNA 15, 1716–1728.

[16] Cavaille, J., Nicoloso, M., and Bachellerie, J. P. (1996) Targeted ribose methylation of RNA in vivo directed by tailored antisense RNA guides, Nature 383, 732–735.

[17] Cavaille, J., and Bachellerie, J. P. (1998) SnoRNA-guided ribose methylation of rRNA: structural features of the guide RNA duplex influencing the extent of the reaction, Nucleic Acids Res 26, 1576–1587.

[18] Kim, S. H., Spensley, M., Choi, S. K., Calixto, C. P., Pendle, A. F., Koroleva, O., Shaw, P. J., and Brown, J. W. (2010) Plant U13 orthologues and orphan snoRNAs identified by RNomics of RNA from Arabidopsis nucleoli, Nucleic Acids Res 38, 3054–3067.

[19] Watkins, N. J., Lemm, I., and Luhrmann, R. (2007) Involvement of nuclear import and export factors in U8 box C/D snoRNP biogenesis, Mol Cell Biol 27, 7018–7027.

[20] Badrock, A. P., Uggenti, C., Wachul, L., Crilly, S., Jenkinson, E. M., Rice, G. I., Kasher, P. R., Lafontaine, D. L., Crow, Y., and O’Keefe, R. T. (2022) Novel intramolecular base - pairing of the U8 snoRNA underlies a Mendelian form of cerebral small vessel disease, bioRxiv.

[21] Boulon, S., Verheggen, C., Jady, B. E., Girard, C., Pescia, C., Paul, C., Ospina, J. K., Kiss, T., Matera, A. G., Bordonne, R., and Bertrand, E. (2004) PHAX and CRM1 are required sequentially to transport U3 snoRNA to nucleoli, Mol Cell 16, 777–787.

[22] Ohno, M., Segref, A., Bachi, A., Wilm, M., and Mattaj, I. W. (2000) PHAX, a mediator of U snRNA nuclear export whose activity is regulated by phosphorylation, Cell 101, 187–198.

[23] Darzacq, X., and Kiss, T. (2000) Processing of intron-encoded box C/D small nucleolar RNAs lacking a 5’,3’-terminal stem structure, Mol Cell Biol 20, 4522–4531.

[24] Jenkinson, E. M., Rodero, M. P., Kasher, P. R., Uggenti, C., Oojageer, A., Goosey, L. C., Rose, Y., Kershaw, C. J., Urquhart, J. E., Williams, S. G., Bhaskar, S. S., O’Sullivan, J., Baerlocher, G. M., Haubitz, M., Aubert, G., Baranano, K. W., Barnicoat, A. J., Battini, R., Berger, A., Blair, E. M., Brunstrom-Hernandez, J. E., Buckard, J. A., Cassiman, D. M., Caumes, R., Cordelli, D. M., De Waele, L. M., Fay, A. J., Ferreira, P., Fletcher, N. A., Fryer, A. E., Goel, H., Hemingway, C. A., Henneke, M., Hughes, I., Jefferson, R. J., Kumar, R., Lagae, L., Landrieu, P. G., Lourenco, C. M., Malpas, T. J., Mehta, S. G., Metz, I., Naidu, S., Ounap, K., Panzer, A., Prabhakar, P., Quaghebeur, G., Schiffmann, R., Sherr, E. H., Sinnathuray, K. R., Soh, C., Stewart, H. S., Stone, J., Van Esch, H., Van Mol, C. E., Vanderver, A., Wakeling, E. L., Whitney, A., Pavitt, G. D., Griffiths-Jones, S., Rice, G. I., Revy, P., van der Knaap, M. S., Livingston, J. H., O’Keefe, R. T., and Crow, Y. J. (2016) Mutations in SNORD118 cause the cerebral microangiopathy leukoencephalopathy with calcifications and cysts, Nat Genet 48, 1185–1192.

[25] Badrock, A. P., Uggenti, C., Wacheul, L., Crilly, S., Jenkinson, E. M., Rice, G. I., Kasher, P. R., Lafontaine, D. L. J., Crow, Y. J., and O’Keefe, R. T. (2020) Analysis of U8 snoRNA Variants in Zebrafish Reveals How Bi-allelic Variants Cause Leukoencephalopathy with Calcifications and Cysts, Am J Hum Genet 106, 694–706.

[26] Huang, G. M., Jarmolowski, A., Struck, J. C., and Fournier, M. J. (1992) Accumulation of U14 small nuclear RNA in Saccharomyces cerevisiae requires box C, box D, and a 5’, 3’ terminal stem, Mol Cell Biol 12, 4456–4463.

[27] Peculis, B. A., and Steitz, J. A. (1994) Sequence and structural elements critical for U8 snRNP function in Xenopus oocytes are evolutionarily conserved, Genes Dev 8, 2241–2255.

[28] Krogh, N., Jansson, M. D., Hafner, S. J., Tehler, D., Birkedal, U., Christensen-Dalsgaard, M., Lund, A. H., and Nielsen, H. (2016) Profiling of 2’-O-Me in human rRNA reveals a subset of fractionally modified positions and provides evidence for ribosome heterogeneity, Nucleic Acids Res 44, 7884–7895.

[29] Cavaille, J., Hadjiolov, A. A., and Bachellerie, J. P. (1996) Processing of mammalian rRNA precursors at the 3’ end of 18S rRNA. Identification of cis-acting signals suggests the involvement of U13 small nucleolar RNA, Eur J Biochem 242, 206–213.

[30] Hadjiolova, K. V., Normann, A., Cavaille, J., Soupene, E., Mazan, S., Hadjiolov, A. A., and Bachellerie, J. P. (1994) Processing of truncated mouse or human rRNA transcribed from ribosomal minigenes transfected into mouse cells, Mol Cell Biol 14, 4044–4056.

[31] Ideue, T., Hino, K., Kitao, S., Yokoi, T., and Hirose, T. (2009) Efficient oligonucleotide-mediated degradation of nuclear noncoding RNAs in mammalian cultured cells, RNA 15, 1578–1587.

[32] Ito, S., Horikawa, S., Suzuki, T., Kawauchi, H., Tanaka, Y., Suzuki, T., and Suzuki, T. (2014) Human NAT10 is an ATP-dependent RNA acetyltransferase responsible for N4-acetylcytidine formation in 18 S ribosomal RNA (rRNA), J Biol Chem 289, 35724–35730.

[33] Ito, S., Akamatsu, Y., Noma, A., Kimura, S., Miyauchi, K., Ikeuchi, Y., Suzuki, T., and Suzuki, T. (2014) A single acetylation of 18 S rRNA is essential for biogenesis of the small ribosomal subunit in Saccharomyces cerevisiae, J Biol Chem 289, 26201–26212.

[34] Bortolin, M. L., Ganot, P., and Kiss, T. (1999) Elements essential for accumulation and function of small nucleolar RNAs directing site-specific pseudouridylation of ribosomal RNAs, EMBO J 18, 457–469.

[35] Czekay, D. P., and Kothe, U. (2021) H/ACA Small Ribonucleoproteins: Structural and Functional Comparison Between Archaea and Eukaryotes, Front Microbiol 12, 654370.

[36] Ganot, P., Caizergues-Ferrer, M., and Kiss, T. (1997) The family of box ACA small nucleolar RNAs is defined by an evolutionarily conserved secondary structure and ubiquitous sequence elements essential for RNA accumulation, Genes Dev 11, 941–956.

[37] Liu, B., and Fournier, M. J. (2004) Interference probing of rRNA with snoRNPs: a novel approach for functional mapping of RNA in vivo, RNA 10, 1130–1141.

[38] Liu, B., Liang, X. H., Piekna-Przybylska, D., Liu, Q., and Fournier, M. J. (2008) Mis-targeted methylation in rRNA can severely impair ribosome synthesis and activity, RNA Biol 5, 249–254.

[39] Karijolich, J., and Yu, Y. T. (2011) Converting nonsense codons into sense codons by targeted pseudouridylation, Nature 474, 395–398.

[40] Thalalla Gamage, S., Sas-Chen, A., Schwartz, S., and Meier, J. L. (2021) Quantitative nucleotide resolution profiling of RNA cytidine acetylation by ac4C-seq, Nat Protoc 16, 2286–2307.

[41] Baserga, S. J., Yang, X. D., and Steitz, J. A. (1991) An intact Box C sequence in the U3 snRNA is required for binding of fibrillarin, the protein common to the major family of nucleolar snRNPs, EMBO J 10, 2645–2651.

[42] Watkins, N. J., Dickmanns, A., and Luhrmann, R. (2002) Conserved stem II of the box C/D motif is essential for nucleolar localization and is required, along with the 15.5K protein, for the hierarchical assembly of the box C/D snoRNP, Mol Cell Biol 22, 8342–8352.

[43] Go, C. D., Knight, J. D. R., Rajasekharan, A., Rathod, B., Hesketh, G. G., Abe, K. T., Youn, J. Y., Samavarchi-Tehrani, P., Zhang, H., Zhu, L. Y., Popiel, E., Lambert, J. P., Coyaud, E., Cheung, S. W. T., Rajendran, D., Wong, C. J., Antonicka, H., Pelletier, L., Palazzo, A. F., Shoubridge, E. A., Raught, B., and Gingras, A. C. (2021) A proximity-dependent biotinylation map of a human cell, Nature 595, 120–124.

[44] Arango, D., Sturgill, D., Alhusaini, N., Dillman, A. A., Sweet, T. J., Hanson, G., Hosogane, M., Sinclair, W. R., Nanan, K. K., Mandler, M. D., Fox, S. D., Zengeya, T. T., Andresson, T., Meier, J. L., Coller, J., and Oberdoerffer, S. (2018) Acetylation of Cytidine in mRNA Promotes Translation Efficiency, Cell 175, 1872–1886 e1824.

[45] Bachellerie, J. P., and Cavaille, J. (1997) Guiding ribose methylation of rRNA, Trends Biochem Sci 22, 257–261.

[46] Dudnakova, T., Dunn-Davies, H., Peters, R., and Tollervey, D. (2018) Mapping targets for small nucleolar RNAs in yeast, Wellcome Open Res 3, 120.

