## Supplementary Information for "Antisense pairing and SNORD13 structure guide RNA cytidine acetylation"

### Table of Contents

| | Page |
|---|---|
| Supplementary Figures S1-S6 | S2-6 |
| Experimental Methods | S7-9 |
| SNORD13 wild-type (WT) and mutant sequences | S10 |
| H45-ITS1 WT and mutant sequences | S11 |
| References | S12 |

### Supplementary Figures

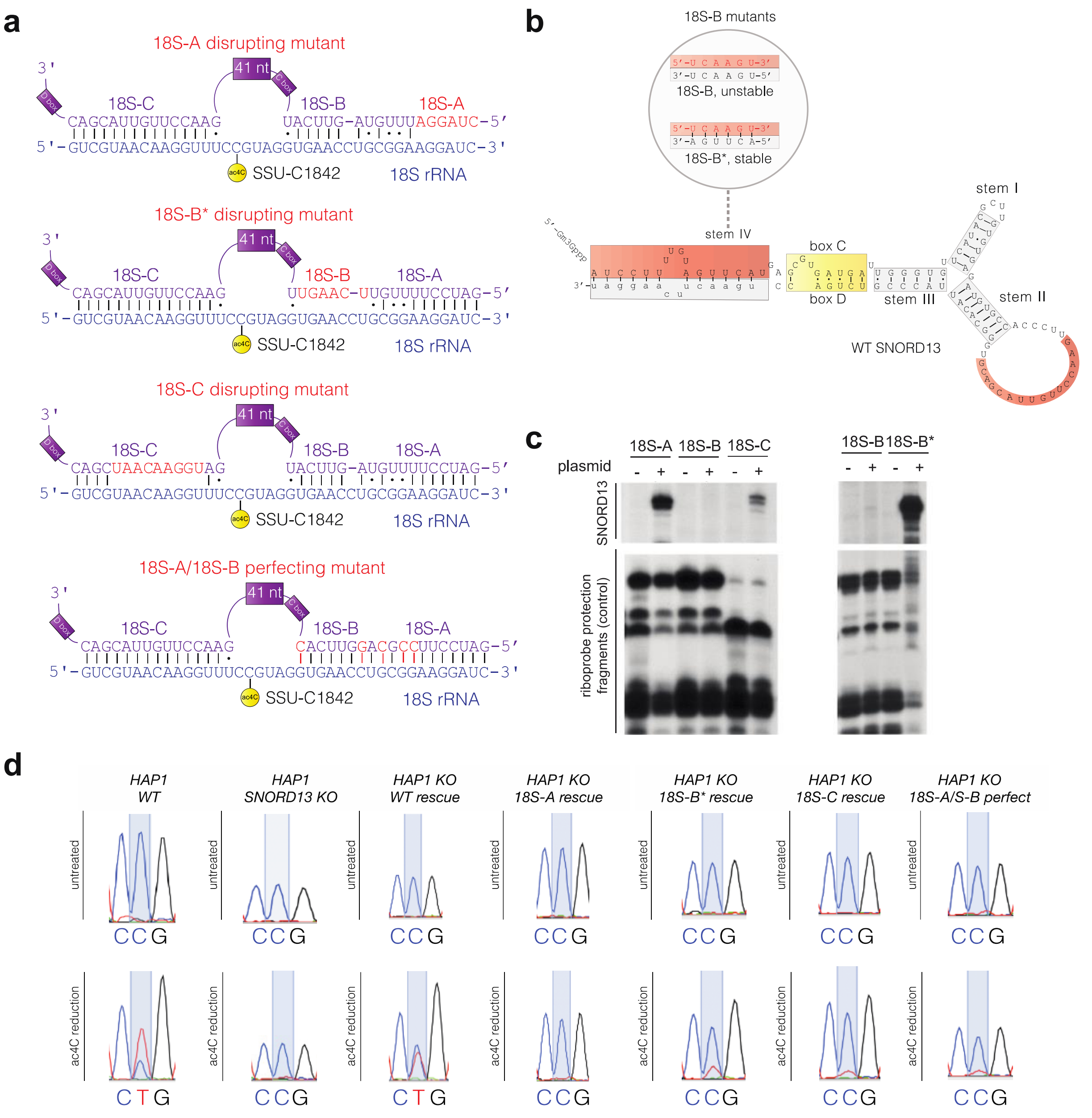


**Figure S1.** (a) Sequences of human SNORD13 mutants analyzed and referenced in Figure 4b. In addition to wild-type ("WT") sequence, disruptive mutations (18S-A, 18S-B*, 18S-C mutants), and a mutant with increased complementarity ("18S-A/18S-B perfect") were explored for rescue of ac$^{4}$C in SNORD13 KO cells. (b) Sequences of SNORD13 18S-B mutants referenced in Figure 4c. Mutation of SNORD13 in mutant '18S-B' disrupts stem IV. Re-introduction of complementarity in construct 18-B* (bottom) allows accumulation and testing of function. (c) Verification of expression of 18S-A, 18S-B, 18S-C, 18S-B* constructs by RNAse A/T1 mapping. Expression of 18S-A/B perfect was verified by Northern (data not shown). (d) Exemplary sequencing traces for WT, SNORD13 KO, and ectopically-expressed SNORD13 rescue experiments referenced in Figure 4d.

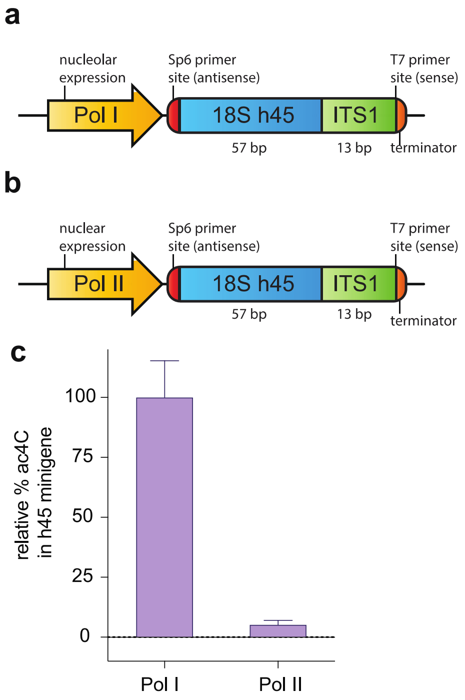


**Figure S2.** (a) Expression schematic for h45-ITS1 minigene construct driven by Pol I and (b) Pol II promoters. The presence of T7 and Sp6 primer binding sites allow differentiation of the h45-ITS1 minigene construct from endogenous helix 45 rRNA in $ac^4C$ sequencing experiments. (c) Bar graph of $ac^4C$ levels at site corresponding to SSU-1842 in h45-ITS1 minigenes driven by Pol I or Pol II promoters, respectively. Values represent $ac^4C$-dependent misincorporation rates normalized relative to the WT Pol1 h45-ITS1, which was set to equal 100%.

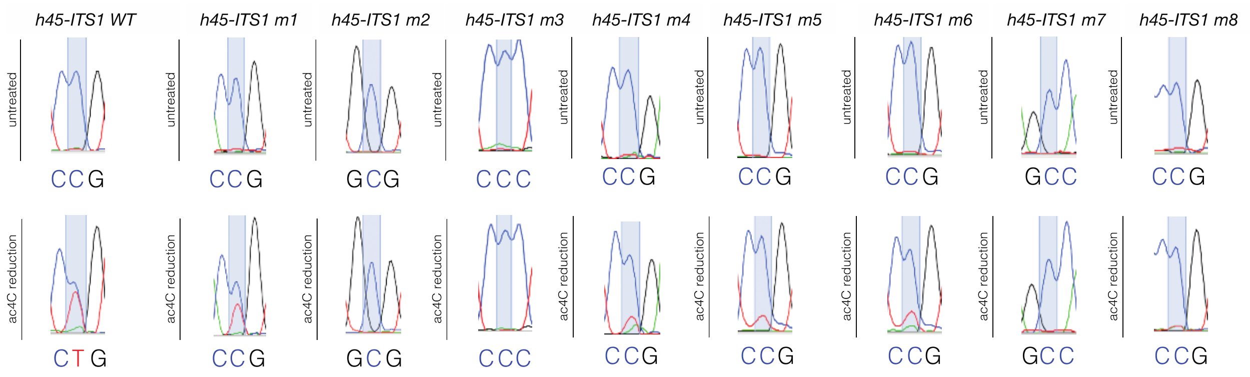


**Figure S3.** Exemplary sequencing traces for modification ectopically-expressed h45-ITS1 minigene substrates referenced in Figure 5b and 5e.

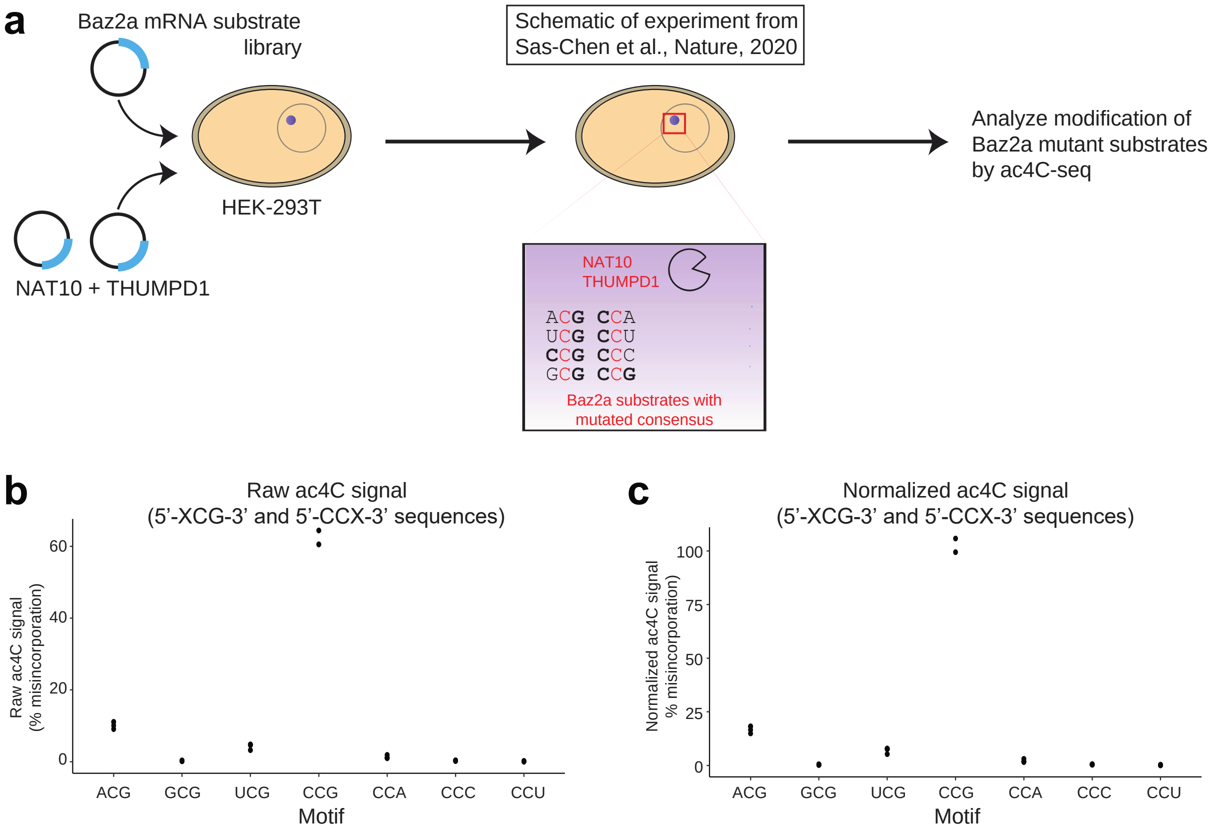


**Figure S4.** Re-analysis of modification of a degenerate RNA substrate library upon overexpression of human NAT10/THUMPD1. (a) Schematic of experiment analyzed from Sas-Chen et al. 2020.[1] HEK-293T cells were co-transfected with NAT10, THUMPD1, and a plasmid expressing a portion of the coding sequence of Baz2a, an mRNA continaing a 5'-CCG-3' sequence that becomes acetylated when NAT10/THUMPD1 are massively overexpressed in this cell line. RNA is isolated and subjected to ac$^4$C-seq to analyze the induction of ac$^4$C at variants of the 5'-CCG-3' consensus sequence. (b) Raw ac$^4$C signal at 5' (XCG) and 3' (CCX) consensus sequences in Baz2a substrate. % misincorporation represents value relative to controls that were not treated with $NaCNBH_3$/HCl. Typical misincorporation rates in untreated controls were 0.1-0.3%. (c) Normalized ac$^4$C signal at 5' (XCG) and 3' (CCX) consensus sequences. 5'-CCG-3' was set to 100%. $NaCNBH_3$/HCl-dependent misincorporation at 5'-ACG-3' and 5'-UCG-3' is consistent with the ability of overexpressed eukaryotic NAT10 to modify these substrates, albeit with greatly reduced eficiency relative to 5'-CCG-3'.

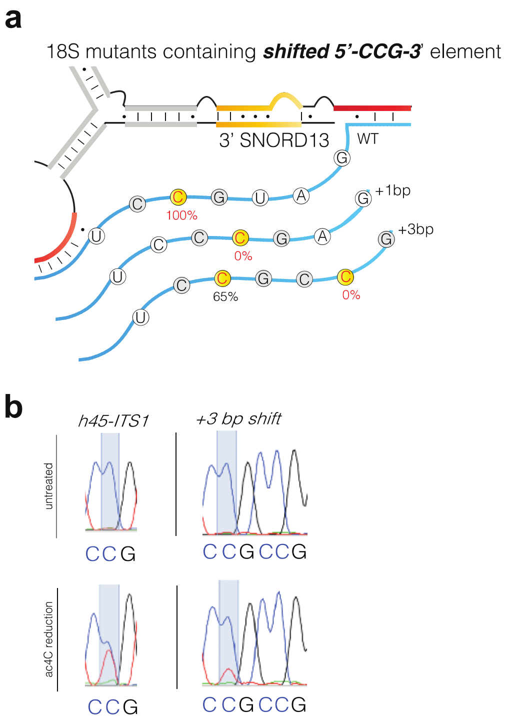


**Figure S5.** (a) Structure of h45-ITS1 substrates in which the 5'-CCG-3' consensus sequence is shifted 1 bp (+1bp) or 3 bp (+3bp; reproduced from Figure 5h). (b) Exemplary sequencing traces showing site of modification in h45-ITS1 substrates containing multiple 5'-CCG-3' sequences in acetylation loop.

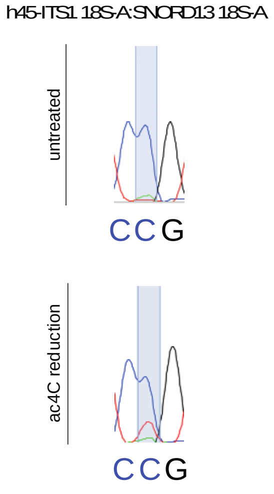


**Figure S6.** Exemplary sequencing traces for SNORD13 18S-A mutant and h45-ITS1 18S-A mutant pair.

### Experimental methods

#### Cell culture, transfection and RNA extraction

HEK-293T and L929 cells were grown in Dulbecco's modified Eagle medium (DMEM; 4.5 g/L glucose). HAP1 cells were grown in Iscove's Modified Dulbecco's Medium (IMDM; 4.5 g/L glucose). All cell culture media were supplemented with 10% fetal bovine serum (PAN biotech), 1 mM Sodium Pyruvate (Gibco) and 1% penicillin/streptomycin (Sigma-Aldrich). Cells were incubated at 37 °C with 5% $CO_2$. L929 and HEK-293T cells (70-80% confluency in 6-well plates) were transiently transfected with plasmids using Lipofectamine 2000 (Invitrogen) according to the manufacturer's instructions. Briefly, 5 µL of Lipofectamine 2000 was diluted in 250 µL Opti-MEM I medium (solution A) and incubated for 5 min. at room temperature. A solution B was prepared with 4 µg of plasmid diluted in 250 µL Opti-MEM I. Solutions A and B were mixed gently (drop-by-drop) and incubated for 20 min. at room temperature. Plasmid-Lipofectamine complexes (500 µL) were then added to seeded L929 or HEK-293T cells. Using GenePulser Xcell (Biorad; 270V, 950µF, 4 mm cuvette), half a T175 culture flask containing SNORD13-KO HAP1 cells at 70-80% confluency was co-electroporated with 8 µg of SNORD13-expressing plasmids and 1 µg of GFP-expressing plasmid (pGFPmax). Forty-eight hours post-transfection, GFP positive cells (~1 million) were sorted using fluorescence-activated cell sorting (BD Influx System (USB) and total RNA was prepared using Tri-reagent (Euromedex) according to the manufacturer's instructions before treatment with RQ1 DNase-RNase free (1u/µl; Promega) and proteinase K (25 mg/mL; Sigma).

#### Preparation of SNORD13 constructs, SNORD13 mutagenesis and RNAse A/T1 protection

A 1.2 kb-long genomic DNA fragment overlapping human SNORD13 gene (hg38: Chr8: 33,512,975-33,514,174) was PCR amplified and cloned into the EcoRI and XbaI sites of the pKS plasmid. SNORD13 mutants were then generated by multi-step PCR (SNORD13 mutant sequences and primer sequences are provided below and in Supplementary Table S1). DNA sequencing was used to confirm the introduced mutations. Plasmids were transfected in L929 cells and total RNA was analyzed by RNAse A/T1 mapping. Briefly, 10 µg of total RNA was hybridized overnight at 42°C with ~ 50 000 cpm of gel-purified [$\alpha^{32}$P]-labeled riboprobes in 10 µL of S1 solution (40 mM PIPES, 400mM NaCl, 1 mM EDTA, 80% formamide). Samples were treated for 60 min at room temperature with 100 µL of RNAse A/T1 buffer (10 mM Tris-Cl pH 7.5, 200 mM NaCl, 5 mM EDTA, 100 mM LiCl, 20-40µg/mL RNase A (Sigma), RNase T1 100 u/mL (Ambion)) followed by a 15-min treatment at 37°C with proteinase K (25 mg/mL, Sigma). After extraction with phenol chloroform-isoamyl alcohol, RNA was precipitated and fractionated by electrophoresis on a 6% acrylamide, 7 M urea denaturing gel. SNORD13-expressing plasmids were used to produce PCR templates for producing antisense riboprobe via T7 in vitro transcription.

**Ac4C-sequencing of endogenous SSU-C1842**

DNase treated Total RNA samples (1 µg) were treated with sodium cyanoborohydride (100 mM in $H_2O$) or vehicle ($H_2O$) in a final reaction volume of 100 µL. Reactions were initiated by the addition of 1 M HCl to a final concentration of 100 mM and incubated for 20 min at room temperature. Reactions were stopped by the addition of 30 µL 1 M tris−HCl pH 8.0. The quenched reactions were adjusted to 200 µL with $H_2O$ and purified via ethanol precipitation. The pelleted RNA was dried using a Speedvac, resuspended in dd$H_2O$, and quantified using a Nanodrop 2000 spectrophotometer. For the reverse transcription reaction with the SuperScript III enzyme, first ~500 pg of treated RNA were incubated with 4.0 pmole of the h45 reverse primer (5'-TAATGATCCTTCCGCAGGTTCACCTAC-3') in 1X Superscript III buffer at 65 °C for 5 min and transferred to ice for 1 min to facilitate annealing. After annealing, reverse transcriptions were performed by adding 200 units of the SuperScript III enzyme, 5 mM DTT, 25 units of RNasin, 500 µM dNTPs (5 mM GTP, 10 mM CTP, ATP, and TTP) and incubating at 55 °C for 60 min. Reactions were quenched by increasing the temperature to 70 °C for 15 min. The cDNA products from the reactions and controls were directly used in PCR. PCR reactions were set up with 2 µL cDNA in 50 µL PCR reaction with Phusion Hot start flex (New England Biolabs). Reaction conditions: 1X supplied HF buffer, 200 µM each dNTP, 2.5 pmol each forward (5'-CGTCGCTACTACCGATTGGATGG-3') and reverse (5'-TAATGATCCTTCCGCAGGTTCACCTAC-3') primers, 2 units of Phusion hot start enzyme, 2 µL template (Thermocycling conditions: 67 °C annealing, 34 cycles). PCR products were run on a 2% agarose gel, stained with SYBR safe, and visualized on a UV transilluminator at 302 nm. Bands of the desired size were excised from the gel. DNA was extracted using a QIAquick gel extraction kit from Zymo and submitted for Sanger sequencing (GeneWiz) using 5 µM of the forward PCR primer. Processed sequencing traces were viewed using 4Peaks software. Peak height for each base was measured, and the percent misincorporation was determined using the equation: “percent misincorporation = (peak intensity of T)/ (sum of C and T base peaks) X 100%”. Misincorporation values were determined by subtracting the average background water control misincorporation levels from that of the corresponding reactions. Then the misincorporation values of each construct were normalized to empty vector value followed by the wild type value.

**Preparation of h45-ITS1 minigene constructs**

A PCR amplified fragment corresponding to the human 18S (57 nt)-ITS1 (13 nt) junction was cloned into the Hind III and BamHI sites of the human ribosomal RNA (Pol-I) minigene phPol1EX (doi/10.1261/rna.1657609) or pcDNA 3.1 which drives expression from a RNA Pol-II (CMV) promoter. Mutated versions were generated by multi-step PCR and DNA sequencing was used to confirm mutations (h45-ITS1 and primer sequences are provided below and in Supplementary Table S1).

**Ac4C-sequencing of h45-ITS1 constructs**

DNase treated Total RNA samples (~5 µg) were treated with sodium cyanoborohydride (100 mM in $H_2O$) or vehicle ($H_2O$) in a final reaction volume of 100 µL. Reactions were initiated by the addition of 1 M HCl to a final concentration of 100 mM and incubated for 20 min at room temperature. Reactions were stopped by the addition of 30 µL 1 M tris−HCl pH 8.0. The quenched reactions were adjusted to 200 µL with $H_2O$ and purified via ethanol precipitation. The pelleted RNA was dried using a Speedvac, resuspended in dd$H_2O$, and quantified using a Nanodrop 2000 spectrophotometer. For the reverse transcription reaction with the SuperScript III enzyme, first ~2 µg of treated RNA were incubated with 4.0 pmole of the T7 reverse primer (5'-TAATACGACTCACTATAG -3') in 1X Superscript III buffer at 65 °C for 5 min and transferred to ice for 1 min to facilitate annealing. After annealing, reverse transcriptions were performed by adding 200 units of the SuperScript III enzyme, 5 mM DTT, 25 units of RNasin, 500 µM dNTPs (5 mM GTP, 10 mM CTP, ATP, and TTP) and incubating at 55 °C for 60 min. Reactions were quenched by increasing the temperature to 70 °C for 15 min. The cDNA products from the reactions and controls were directly used in PCR. PCR reactions were set up with 2 µL cDNA in 50 µL PCR reaction with Phusion Hot start flex (New England Biolabs). Reaction conditions: 1X supplied HF buffer, 2.5 pmol each SP6 forward (5'-ATTTAGGTGACACTATAGAA -3') and T7 reverse (5'-TAATACGACTCACTATAG-3') primers, 200 µM each dNTP, 2 units of Phusion hot start enzyme, 2 µL template (Thermocycling conditions: 52 °C annealing, 33 cycles). PCR products were run on a 2% agarose gel, stained with SYBR safe, and visualized on a UV transilluminator at 302 nm. Bands of the desired size were excised from the gel. DNA was extracted using a QIAquick gel extraction kit from Zymo and submitted for Sanger sequencing (GeneWiz) using 5 µM of the SP6 forward PCR primer. Processed sequencing traces were viewed using 4Peaks software. Peak height for each base was measured, and the percent misincorporation was determined using the equation: “percent misincorporation = (peak intensity of T)/ (sum of C and T base peaks) X 100%”. Misincorporation values were determined by subtracting the average background water control misincorporation levels from that of the corresponding reactions. Then the misincorporation values of each construct were normalized to the wild type value.

### SNORD13 WT and mutant sequences

```
>U13-wt
GATCCTTTTGTAGTTCATGAGCGTGATGATTGGGTGTTCATACGCTTGTGTGAGATGTGCCACCCTTGAACCTTG
TTACGACGTGGGCACATTACCCGTCTGACC

>U13-Cmut
GATCCTTTTGTAGTTCATGAGCGACTACTTTGGGTGTTCATACGCTTGTGTGAGATGTGCCACCCTTGAACCTTG
TTACGACGTGGGCACATTACCCGTCTGACC

>U13-Dmut
GATCCTTTTGTAGTTCATGAGCGTGATGATTGGGTGTTCATACGCTTGTGTGAGATGTGCCACCCTTGAACCTTG
TTACGACGTGGGCACATTACCCGTGACTCC

>U13-stem-I
GATCCTTTTGTAGTTCATGAGCGTGATGATTGGGTGTTCATACGCTTGTCACTGATGTGCCACCCTTGAACCTTG
TTACGACGTGGGCACATTACCCGTCTGACC

>U13-stem-I-comp
GATCCTTTTGTAGTTCATGAGCGTGATGATTGGGTGTAGTGACGCTTGTCACTGATGTGCCACCCTTGAACCTTG
TTACGACGTGGGCACATTACCCGTCTGACC

>U13-stem-II
GATCCTTTTGTAGTTCATGAGCGTGATGATTGGGTGTTCATACGCTTGTGTGAGATGTGCCACCCTTGAACCTTG
TTACGACGTGGCGTGTTTACCCGTCTGACC

>U13-stem-II-comp
GATCCTTTTGTAGTTCATGAGCGTGATGATTGGGTGTTCATACGCTTGTGTGAGAACACGCACCCTTGAACCTTG
TTACGACGTGGCGTGTTTACCCGTCTGACC

>U13-stem-III
GATCCTTTTGTAGTTCATGAGCGTGATGATTCCCAGTTCATACGCTTGTGTGAGATGTGCCACCCTTGAACCTTG
TTACGACGTGGGCACATTACCCGTCTGACC

>U13-stem-III-comp
GATCCTTTTGTAGTTCATGAGCGTGATGATTCCCAGTTCATACGCTTGTGTGAGATGTGCCACCCTTGAACCTTG
TTACGACGTGGGCACATTTGGGGTCTGACC

>U13-stem-IV
GATCCTTTTGTAGTTCATGAGCGTGATGATTGGGTGTTCATACGCTTGTGTGAGATGTGCCACCCTTGAACCTTG
TTACGACGTGGCACATTACCCGTCTGACCACTTGATCAAGGATC

>U13-stem-IV-comp
GATCCTTTTGTTCAAGTTGAGCGTGATGATTGGGTGTTCATACGCTTGTGTGAGATGTGCCACCCTTGAACCTTG
TTACGACGTGGGCACATTACCCGTCTGACCACTTGATCAAGGATC

>U13-18S-A
CTAGGAATTGTAGTTCATGAGCGTGATGATTGGGTGTTCATACGCTTGTGTGAGATGTGCCACCCTTGAACCTTG
TTACGACGTGGGCACATTACCCGTCTGACC

>U13-18S-B
GATCCTTTTGTACAAGTAGAGCGTGATGATTGGGTGTTCATACGCTTGTGTGAGATGTGCCACCCTTGAACCTTG
TTACGACGTGGGCACATTACCCGTCTGACC

>U13-18S-C
ATCCTTTTGTAGTTCATGAGCGTGATGATTGGGTGTTCATACGCTTGTGTGAGATGTGCCACCCTTGATGGAACA
ATCGACGTGGGCACATTACCCGTCTGACC

>hU13-18S_AB perfect comp
GATCCTTCCGCAGGTTCACGAGCGTGATGATTGGGTGTTCATACGCTTGTGTGAGATGTGCCACCCTTGAACCTT
GTTACGACGTGGGCACATTACCCGTCTGACCGTGAACCTGCGGAAGGATC
```

### h45-ITS1 WT and mutant sequences

**>h45-WT**
AGAGGAAGTAAAAGTCGTAACAAGGTTTCCGTAGGTGAACCTGCGGAAGGATCATTAacggagcccggag
**>h45_m1**
AGAGGAAGTAAAAGTCGTAACAAGGTTACCGTAGGTGAACCTGCGGAAGGATCATTAacggagcccggag
**>h45_m2**
AGAGGAAGTAAAAGTCGTAACAAGGTTTGCGTAGGTGAACCTGCGGAAGGATCATTAacggagcccggag
**>h45_m3**
AGAGGAAGTAAAAGTCGTAACAAGGTTTCCCTAGGTGAACCTGCGGAAGGATCATTAacggagcccggag
**>h45_m4**
AGAGGAAGTAAAAGTCGTAACAAGGTTTCCGAAGGTGAACCTGCGGAAGGATCATTAacggagcccggag
**>h45_m5**
AGAGGAAGTAAAAGTCGTAACAAGGTTTCCGTTGGTGAACCTGCGGAAGGATCATTAacggagcccggag
**>h45_m6**
AGAGGAAGTAAAAGTCGTAACAAGGTTTCCGTACGTGAACCTGCGGAAGGATCATTAacggagcccggag
**>h45_m7**
AGAGGAAGTAAAAGTCGTAACAAGGTTAGCCATCGTGAACCTGCGGAAGGATCATTAacggagcccggag
**>h45_m8**
AGAGGAAGTAAAAGTCGTAACAAGGTTCCCGTAGATGAACCTACGGAAGGATCATTAacggagcccggag
**>h45-ACG**
AGAGGAAGTAAAAGTCGTAACAAGGTTTACGTAGGTGAACCTGCGGAAGGATCATTAacggagcccggag
**>h45-UCG**
AGAGGAAGTAAAAGTCGTAACAAGGTTTUCGTAGGTGAACCTGCGGAAGGATCATTAacggagcccggag
**>h45-CCA**
AGAGGAAGTAAAAGTCGTAACAAGGTTTCCATAGGTGAACCTGCGGAAGGATCATTAacggagcccggag
**>h45-CCU**
AGAGGAAGTAAAAGTCGTAACAAGGTTTCCUTAGGTGAACCTGCGGAAGGATCATTAacggagcccggag
**>h45_ins1**
AGAGGAAGTAAAAGTCGTAACAAGGTTTCCCGTAGGTGAACCTGCGGAAGGATCATTAacggagcccggag
**>h45_ins2**
AGAGGAAGTAAAAGTCGTAACAAGGTTTCCCCGTAGGTGAACCTGCGGAAGGATCATTAacggagcccggag
**>h45_ins3**
AGAGGAAGTAAAAGTCGTAACAAGGTTTCCGTACGGTGAACCTGCGGAAGGATCATTAacggagcccggag
**>h45_ins4**
AGAGGAAGTAAAAGTCGTAACAAGGTTTCCGTACCGGTGAACCTGCGGAAGGATCATTAacggagcccggag
**>h45_ins5**
AGAGGAAGTAAAAGTCGTAACAAGGTTTCCCGTACGGTGAACCTGCGGAAGGATCATTAacggagcccggag
**>h45_ins6**
AGAGGAAGTAAAAGTCGTAACAAGGTTTCCCGTACCGGTGAACCTGCGGAAGGATCATTAacggagcccggag
**>shifted 5′-CCG-3′ +1**
AGAGGAAGTAAAAGTCGTAACAAGGTTTCCCGAGGTGAACCTGCGGAAGGATCATTAacggagcccggag
**>shifted 5′-CCG-3′ +3**
AGAGGAAGTAAAAGTCGTAACAAGGTTTCCGCCGGTGAACCTGCGGAAGGATCATTAacggagcccggag
**>h45_18SA**
AGAGGAAGTAAAAGTCGTAACAAGGTTTCCGTAGGTGAACCTGCGGTTCCTACATTAacggagcccggag

### References

[1] Sas-Chen, A., Thomas, J. M., Matzov, D., Taoka, M., Nance, K. D., Nir, R., Bryson, K. M., Shachar, R., Liman, G. L. S., Burkhart, B. W., Gamage, S. T., Nobe, Y., Briney, C. A., Levy, M. J., Fuchs, R. T., Robb, G. B., Hartmann, J., Sharma, S., Lin, Q., Florens, L., Washburn, M. P., Isobe, T., Santangelo, T. J., Shalev-Benami, M., Meier, J. L., and Schwartz, S. (2020) Dynamic RNA acetylation revealed by quantitative cross-evolutionary mapping, *Nature 583*, 638-643.
